# Chronic alcohol exposure induces cerebral microbleeds by TGF-β1/Smad signaling pathway mediated remodeling of cerebral small vessels

**DOI:** 10.1101/2024.02.07.579406

**Authors:** Hengjian Lu, Hongxuan Wang, Xiangpen Li, Xinrou Lin, Chenguang Li, Wanru Chen, Lubin Zou, Jingrui Pan, Xiaoni Zhang, Lei He, Xiaoming Rong, Ying Peng

## Abstract

**Background:** Long-term heavy drinking is a major risk factor for cerebral microbleeds(CMBs), which are increasingly gaining attention as a pathological phenotype of cerebral small vessel diseases(CSVD). Under pathological conditions, remodeling of the extracellular matrix(ECM) on the walls of small vessels causes disarray in the structure and function of these vessels, leading to cerebral small vessel sclerosis and consequent rupture and bleeding. This can result in cognitive and emotional disorders, abnormal gait and increased risk of falling. However, the mechanisms underlying how long-term alcohol consumption leads to CMBs and decline in motor function remain unknown.

**Methods:** We constructed a chronic alcohol exposure mouse model and measured the deposition of ECMs on the small vessels in motor-related brain regions. The presence of microbleeds was confirmed through Prussian blue staining and Magnetic Resonance Imaging. We also extracted primary cerebral microvascular smooth muscle cells (CMVSMCs) from the newborn mice and explored the effects of alcohol on the phenotypic transformation and substance synthesis function. Additionally, we conducted interventional experiments on the cell and animal models with an anti-fibrotic drugs Pirfenidone(PFD).

**Results:** We found that mice with long-term alcohol exposure showed decreased motor function. In their motor-related brain regions, such as the motor cortex(MC), thalamus/basal ganglia(Tha/BG), and cerebellum(CB), we observed microbleeds. On the small vessels in these areas, we detected excessive deposited ECM proteins. In vitro experiments with primary CMVSMCs revealed that after alcohol treatment, the cells underwent a transformation into fibroblast-like phenotypes, and excessive production of the aforementioned ECM proteins, which is regulated by upstream TGFβ1/Smad signaling pathway. Additionally, PFD applied on cell and animal models could reverse the above processes to some extent.

**Conclusions:** Our study found that the remodeling of ECM accompanied by activation of TGF-β1/Smad signaling pathway may be involved in alcohol-induced CMBs. It could be a potential therapeutic target for CMBs or CSVD.

## 1. Introduction

Alcohol, as a long-standing psychoactive substance in human society, is widely consumed globally and have significant effects on various physiological systems^1^. The World Health Organization (WHO) points out that harmful use of alcohol is a cause of more than 200 diseases and injuries^2^. Alcohol can raise blood pressure and increase the risk of stroke, offering no protective effect on cerebral blood vessels^3^.

CMBs are a typical radiological manifestation of CSVD, characterized by focal hemosiderin deposits around blood vessels. In MRI blood-sensitive sequences such as T2*-GRE or SWI, they appear as circular or oval lesions with no signal or extremely low signal, generally less than 10mm in diameter^4,5^.With the widespread application of high-field strength MRI and techniques, the detection rate of microbleeds has gradually increased. Microbleeds are more common in diseases such as Alzheimer’s, cerebral amyloid angiopathy, stroke, and brain trauma^6^.

Some epidemiological studies suggest that alcohol consumption is an independent risk factor for CMBs independent of hypertension and diabetes, particularly heavy drinking. And heavy drinking is closely clinically associated with deep CMBs.^7–9^.

However, there is currently no research on the mechanisms linking chronic alcohol consumption and CMBs. Therefore, investigating whether chronic alcohol consumption can cause CMBs and the underlying mechanisms holds significance.

Cerebral small vessel arteriosclerosis is the most common pathological subtype in CSVD, characterized by pathological thickening of the small artery walls and abnormal deposition of ECM proteins. This increases the fragility of the small vessels, leading to vessel rupture^10–12^. In a rat model of CSVD induced by hypertension, abnormal deposition of ECM proteins such as collagen I, collagen IV, fibronectin and laminin on the cerebral small vessel walls can be observed, indicating changes in the composition of the cerebral small vessel walls^13,14^. It is still unclear whether alcohol causes abnormal deposition of ECM proteins in the walls of cerebral small vessels.

ECM proteins are primarily synthesized by vascular smooth muscle cells and are maintained in a constantly renewed state^15^. Alcohol can directly affect vascular smooth muscle cells, impairing their physiological functions, including promoting cell apoptosis^16^, damaging ion channel functions, thereby disrupting the contractile and relaxant functions of small vessels, which affects cerebral blood flow^17,18^. However, how these mechanisms affect the occurrence and progression of small vessel disease is not yet clear. And whether alcohol impacts the synthesis of ECM proteins also remains unknown. Under pathological conditions, smooth muscle cells undergo phenotypic transformation^19^, shifting from a quiescent contractile phenotype to a synthetic one, characterized by enhanced proliferation and migration^20^. Decorin and Lumican are markers for the transformation of vascular smooth muscle cells into fibroblast-like cells, a type of synthetic phenotype^21,22^. This phenotypic transformation also results in the overproduction of the ECM. Therefore, whether alcohol promotes this phenotypic transformation in vascular smooth muscle cells is an important intermediate step in vascular remodeling.

For a long time, TGF-β1 has been considered the most important ECM regulatory factor^23^. The TGF-β1/Smad pathway plays an important role in alcohol-induced liver fibrosis. Phosphorylation of Smad2/3 downstream of TGF-β1 signaling and subsequent nuclear translocation initiates the expression of ECM proteins such as collagen and fibronectin^24,25^. Inhibiting the TGF-β/Smad signaling pathway in hepatic stellate cells can suppress alcohol-induced liver fibrosis^26^. In models of CSVD, the TGF-β1/Smad signaling pathway has been confirmed to undergo changes, and it may be involved in mediating the synthesis of ECM proteins^13^. There is relatively little research on the TGF-β1/Smad signaling pathway in alcohol-related cerebral vascular diseases and even in central nervous system damage. Therefore, focusing on this pathway could be a valuable entry point in exploring the mechanisms underlying alcohol-induced CMBs.

Gait abnormalities and the risk of falling are issues of considerable clinical concern. According to some clinical studies, CMBs are associated with gait and balance disorders, and the presence of these microbleeds in motor-related pathway, such as the frontal lobes, basal ganglia and thalamus, can explain this phenomenon^27,28^. Long-term and severe alcohol abuse can have detrimental effects on lower limb function, and with time, it can lead to serious gait ataxia^29^. Therefore, whether the motor function abnormalities caused by alcohol abuse are a result of CMBs is a topic worth exploring.

In this study, we attempt to explore whether chronic alcohol exposure can mediate the remodeling of the ECM of cerebral small vessels through changes in the TGF-β1/Smad signaling pathway, ultimately leading to CMBs and a decline in motor abilities. Additionally, we conducted interventional studies with PFD, a specific inhibitor of TGF-β1, to determine if it could reverse the aforementioned processes.

## 2. Materials and methods

### 2.1. Animals

Adult 6–8-week C57BL/6J male mice (18–25 g) were procured from Yancheng Bioscience, Guangzhou, China. The mice were adaptively bred for 7 days in a standard environment (temperature of 25 ±2℃, 12-h light/12-h dark cycle, with adequate water and food), and then randomly divided into corresponding experimental groups.

### 2.2. Chronic alcohol consumption model

The chronic alcohol consumption model follows the previously established method^30,31^. Briefly, the model group is administered alcohol by gavage (4g/kg, 25% ethanol), while the control group is given an equivalent volume of water, based on the body weight of the mice in the model group, and this is continued for 28 days. During the gavage procedure, animal ethics are strictly followed, ensuring gentle handling to minimize discomfort or coughing in the mice. This model is defined as a binge drinking model and effectively simulates the pattern of long-term excessive alcohol consumption in humans^32^. After 28 days of uninterrupted gavage treatment, the mice are allowed a rest period of 24 hours before being euthanized or used for behavioral experiments.

### 2.3. Pirfenidone administration

Pirfenidone(Selleck, USA) is a specific inhibitor of TGF-β1, capable of inhibiting both the production of TGF-β and the collagen synthesis stimulated by TGF-β^33,34^. In vitro dissolution tests confirmed that pirfenidone can be completely dissolved in a 25% ethanol solution. Therefore, this study used the aforementioned dissolution method to prepare the medication for the drug intervention group (4g/kg/day 25% ethanol w/v + pirfenidone 200mg/kg/day).

### 2.4. Rotarod test

The rotarod test, first proposed by Dunham and Miya, is used to assess motor coordination and balance abilities ^35^. It has been widely used in assessing motor functions in cases of intracranial hemorrhage, multiple sclerosis, and traumatic brain injury^36–38^.Referring to the previous experimental protocols^39^, before the official test begins, let the mice undergo adaptive training on the rotating rod at a constant speed of 15 rpm for 15 minutes. During this period, if a mouse falls off, it will be put back on the rotating rod to continue training until it can walk steadily on it. In the official test phase, the mode changes to acceleration, where the rod speeds up from 4 rpm to 40 rpm within 5 minutes. The exhaustion time is set as one instance; if the mouse falls off the platform, there is no electrical stimulation. Record the duration of the mouse’s movement on the rotating rod via Five-Mouse Universal Rotarod Experiment System(JLBehv-RRTG, shanghai, China); a duration of 5 minutes without falling is recorded as 300 seconds. Each mouse is tested three times with a 15-minute interval between tests. Finally, the average of the three tests is taken for statistical analysis.

### 2.5. Pre-treatment of coronal frozen sections of brain tissue

After anesthetizing the mice in each group with 1.25% ready-to-use tribromoethanol (0.2ml/10g) (2126A, Aibeibio, China) via intraperitoneal injection, the mice were placed in a supine position on the surgical table with their thoracic cavity fully exposed. The right atrial appendage was cut open, and a puncture needle was inserted through the apex of the heart, followed by rapid infusion with approximately 30ml pre-chilled physiological saline. Once the effluent from the right atrial appendage turns clear, the infusion was immediately switched to pre-chilled 4% paraformaldehyde(BL302A, Biosharp, China), using about 30ml. After the perfusion was completed, the mouse skull was cut open to expose the brain tissue, and the entire mouse brain was carefully removed. The brain was then subjected to gradient dehydration with 4% paraformaldehyde solutions containing 15% and 30% sucrose(BS085, Biosharp, China). After dehydration, the brain tissue was embedded in OCT(4583, SAKURA, Japan) and coronal sections were made in a freezing microtome (LEICA CM1950, Germany) at −20°C with a thickness of 20µm. Brain slices containing MC, Tha/BG, and CB were then mounted on slides and naturally air-dried.

### 2.6. Prussian Blue Staining

In CMBs, localized deposits of iron-containing hemosiderin are observed around blood vessels^40^. The principle of staining lies in using potassium ferrocyanide to extract trivalent iron ions from proteins. This results in a reaction that forms an insoluble blue ferric ferrocyanide precipitate^41^. The Prussian Blue staining solution (G1029, Servicebio, China) is prepared as a working solution, and the brain slices are stained according to the protocol provided in the manual. Subsequently, a glass slide scanning imaging system(SQS-40P, SQRAY, China) is used for statistical analysis of the staining, with particular attention paid to the staining in the MC, Tha/BG, and CB. The number and area of positive stains are counted by ImageJ v1.53q, with this task being carried out by two independent researchers.

### 2.7. Immunofluorescence staining (IF)

Before immunofluorescence staining, the sections undergo antigen retrieval. First, the sections are dried at 37 °C for 2 hours, then immersed in boiling 10 mM citrate buffer (AR0024, BOSTER,China), and continuously heated in a microwave for 10 minutes, maintaining the temperature above 92 °C. Then, 5% goat serum(AR0009, BOSTER,China) is added to block the brain sections at room temperature for 1 hour, followed by the addition of diluted primary antibody for overnight incubation at 4 °C. The strategy for dual immunofluorescence staining is as follows: Mouse anti-α-SMA (R&D, MAB1420) were co-stained with rabbit anti-Collagen I (Abmart, T61022), rabbit anti-Collagen IV (Proteintech, 55131-1), rabbit anti-Fibronectin (Proteintech, 15613-1) and rabbit anti-Laminin (Abmart, TD3618). After overnight incubation with the primary antibody, adopt the following secondary antibody incubation strategy: Mix Alexa Fluor 488-labeled Goat Anti-Mouse IgG(H+L)(Beyotime, A0428) and Alexa Fluor 555-labeled Donkey Anti-Rabbit IgG(H+L)(Beyotime, A0453) together and incubated at room temperature for 2 hours. After completing the coverslipping, use Imager D2(Carl Zeiss, Germany) to observe the localization of the microvessels and the deposition of ECM proteins on the vessel walls. For each group, randomly select 2-3 vascular cross-sections from the MC, Tha/BG, and CB region of the three independent samples for analysis.

For immunofluorescence of CMVSMCs, first plate the cells onto a 24-well plate. After specific treatment, fix the cells with 4% paraformaldehyde(BL302A, Biosharp, China) at room temperature for 15 minutes, then permeabilize with 0.5% Triton X-100 for 10 minutes, followed by blocking with 5% goat serum(AR0009, BOSTER,China) for 30 minutes. The primary antibody incubation strategy is as follows: rabbit anti-Collagen I (Proteintech,14695-1-AP), rabbit anti-Collagen IV (Proteintech,55131-1), rabbit anti-Fibronectin (Proteintech,15613-1) and rabbit anti-Laminin (Proteintech,23498-1-AP), rabbit anti-Smad2/3(Immunoway,YT4332), rabbit anti-pSmad2/3(Immunoway,YP0362) were used for single staining, followed by overnight incubation at 4℃. The next day, incubate with the following fluorescent secondary antibodies at room temperature for 1 hour: Alexa Fluor 555-labeled Donkey Anti-Rabbit IgG(H+L)(Beyotime, A0453). After staining the cell nuclei with DAPI (S2110, Solarbio, China), use an inverted microscope(Nikon, N31373) to observe the expression of ECM proteins and the expression and nuclear localization of phosphorylated Smad in each group.

### 2.8. Real-time quantitative PCR

Total RNA is extracted from CMVSMCs in three independent replicate experiments using RNAiso Plus (RC112, Vazyme, China). The mRNA is then reverse transcribed using HiScript III RT SuperMix (R222, Vazyme, China). For RT-PCR analysis, a 20μl system of ChamQ SYBR Master Mix (Low ROX Premixed) (Q331, Vazyme, China) is employed on the ABI QuantStudio5 System(Thermo Fisher, USA). GAPDH is chosen as the internal reference gene. The primer sequences are as follows: COL1A1 forward, 5’-GCTCCTCTTAGGGGCCACT-3’; COL1A1 reverse, 5’-CCACGTCTCACCATTGGGG-3’; COL4A2 forward, 5’-GACCGAGTGCGGTTCAAAG −3’; COL4A2 reverse, 5’-CGCAGGGCACATCCAACTT-3’; FN1 forward, 5’-GCTCAGCAAATCGTGCAGC-3’; FN1 reverse, 5’-CTAGGTAGGTCCGTTCCCACT-3’; LAMB1 forward, 5’-AGACCCGAAGAAAAGACAGGC-3’; LAMB1 reverse, 5’-CCATAGGGCTAGGACACCAAA-3’; DECORIN forward, 5’-TCTTGGGCTGGACCATTTGAA-3’; DECORIN reverse, 5’-CATCGGTAGGGGCACATAGA-3’; LUMICAN forward, 5’-CTCTTGCCTTGGCATTAGTCG-3’; LUMICAN reverse, 5’-GGGGGCAGTTACATTCTGGTG-3’; TGFB1 forward, 5’-CTCCCGTGGCTTCTAGTGC-3’; TGFB1 reverse, 5’-GCCTTAGTTTGGACAGGATCTG-3’; SMAD2 forward, 5’-ATGTCGTCCATCTTGCCATTC-3’; SMAD2 reverse, 5’-AACCGTCCTGTTTTCTTTAGCTT-3’; SMAD3 forward, 5’-CACGCAGAACGTGAACACC-3’; SMAD3 reverse, 5’-GGCAGTAGATAACGTGAGGGA-3’; GAPDH forward, 5′-AGGTCGGTGTGAACGGATTTG-3′; and GAPDH reverse, 5′-TGTAGACCATGTAGTTGAGGTCA-3′.

### 2.9. Western blotting

Homogenize mouse brain tissues(MC, Tha/BG, and CB) and CMVSMCs in ice-cold lysis buffer (RIPA(P0013B, Beyotime, China): PMSF(ST506, Beyotime, China) = 100:1) for 30 minutes, followed by high-speed centrifugation (4°C, 12000rpm, Eppendorf, 5424R) for 20 minutes to collect the protein supernatant. Separate 20μg of total protein using 7.5% polyacrylamide gel electrophoresis (PG211, Epizyme, China), then transfer onto a polyvinylidene fluoride membrane (PR05505, Millipore, Ireland), and subsequently block with 5% skimmed milk(Wandashan, China) at room temperature for 1 hour. After blocking, incubate overnight at 4°C using the following primary antibody incubation protocol: rabbit anti-Collagen I (Proteintech,14695-1-AP), rabbit anti-Collagen IV (Proteintech,55131-1), rabbit anti-Fibronectin (Proteintech,15613-1) and rabbit anti-Laminin (Proteintech,23498-1-AP), rabbit anti-Smad2/3(Immunoway,YT4332), rabbit anti-pSmad2/3(Immunoway,YP0362), mouse anti-TGFβ1(Immunoway,YM4305), goat anti-Decorin(R&D, AF1060),mouse anti-Lumican(R&D, MAB28461) and rabbit anti-β-actin(Proteintech, 81115-1). After the primary antibody incubation, incubate with the corresponding species-specific HRP-conjugated secondary antibody(GAR007, Goat Anti-Rabbit IgG(H+L), GAM007, Goat Anti-Mouse IgG(H+L), Multi Sciences) at room temperature for 2 hours. Finally, detect the signal using ECL enhanced chemiluminescence solution(SQ201L, Epizyme, China) for development and signal detection(E-BLOT, China).

### 2.10. Extraction and Culture of primary cerebral microvascular smooth muscle cells(CMVSMCs)

Euthanize the mouse(1-3 days) by cervical dislocation, then soak it in 75% alcohol for 5 minutes. Move to a laminar flow hood, prone and secure the mouse, cut open the skull, and aseptically remove the brain tissue. Under a dissecting microscope(Phenix, China), separate the cortical tissue, and cut the tissue into 1 mm^3^ pieces, placing them in PBS with double antibiotics(Procell, China). After washing the tissue once with PBS, transfer it to a centrifuge tube and add digestion fluid (collagenase II(Sigma, USA)). Digest on a shaking water bath at 37°C for 1 hour. Gently pipette until no clumps remain in the liquid. Centrifuge the tissue suspension at 800g for 5 minutes, discard the supernatant. Resuspend the sediment in 20% BSA, then centrifuge at 1200g for 25 minutes and discard the supernatant. Resuspend the microvessel segments in PBS, centrifuge at 400g for 5 minutes, discard the supernatant, and retain the sediment. Resuspend the mouse cerebral microvascular smooth muscle cells in complete culture medium(Procell, China) and seed them in a poly-L-lysine(Procell, China) precoated culture dish. Incubate the cells at 37°C in a 5% CO2 incubator without disturbance. After 48 hours, change the culture medium, and subsequently change the medium every 3 days. Once the cells have reached confluency, they are ready for use. Use Rabbit anti-α-SMA(Proteintech, 14395-1-AP) for immunofluorescence to identify the cells.

CMVSMCs were divided into control group, alcohol treatment group (100mM ethanol), and alcohol+PFD group (100mM ethanol + 0.3μg/μl, Pirfenidone). The concentration of ethanol was determined based on cell viability, and the dosage of Pirfenidone was established based on previous research^42,43^. The cells were harvested after 24h of treatments.

### 2.11. Cell viability

Seed CMVSMCs at a density of 3000 cells/well in a 96-well plate. Culture the cells in a CO2 incubator at 37°C for 24 hours, then apply different treatments according to the cell culture grouping. After 12/24/48 hours of treatment, add 10μL/well of CCK8 solution(GLPBIO, USA). Incubate the plate in the incubator for 2 hours, then measure the absorbance at 450 nm using microplate reader(CLARIOstar, Germany). The cell survival rate (%) = [(absorbance of experimental well -absorbance of blank well) / (absorbance of control well -absorbance of blank well)] ×100%. Calculate the cell survival rate for each experimental well and take the average for each group as the survival rate for that treatment group for subsequent statistical analysis.

### 2.12. Morphometric analysis of CMVSMCs

Under pathological conditions, vascular smooth muscle cells can shift from a contractile phenotype to a synthetic phenotype, morphologically transitioning from a “spindle shape” to an “epithelial-like shape.” This is characterized by a reduction in the length of the cell’s long axis and a decrease in branching^44^. In this experiment, we selected representative fields of view for comparison, and to display the cellular morphology more clearly, we converted the images from light microscopy into binary images using ImageJ v1.53q.

### 2.13. Small Animal Magnetic Resonance Imaging

Magnetic resonance imaging was performed using the Southern Medical University small animal MRI (BRUKER, PharmaScan70/16 US, 7.0T). Mice were anesthetized under a flow of oxygen with 0.5-2% isoflurane gas, then stabilized in the device, aligning the brain with the central region of the magnetic field. SWI sequences were used for the detection of microbleeds.

### 2.14. Statistical analysis

All of the data are presented as mean ±standard error of the mean (SEM). For statistical analysis, independent sample t-tests were used for two groups, and one-way ANOVA was used for multiple groups. Before statistical testing, data were assessed for normality and homogeneity of variance. If these criteria were not met, the Kruskal-Wallis H test or Mann-Whitney U test was used. The analysis was uniformly performed using GraphPad Prism 9.5.1 software. A P-value less than 0.05 (two-tailed) was considered statistically significant.

## 3. Results

### 3.1 Chronic alcohol exposure induces CMBs

To investigate whether chronic alcohol exposure can induce microbleeds, mice were administered with either alcohol or water through gavage for a continuous period of 28 days. After the treatment, brain tissue sections were stained with Prussian Blue (Figure.1A). We conducted a statistical analysis of CMBs in the MC, Tha/BG and CB, the regions of the motor circuit. The results showed that in these regions, mice in the alcohol group had a greater number of microbleeds (blue deposits) in their brain tissues compared to the control group(Figure.1B), and the average CMH size were larger(Figure.1C). This suggests that chronic alcohol consumption in mice could directly lead to CMBs.

**Figure.1.**
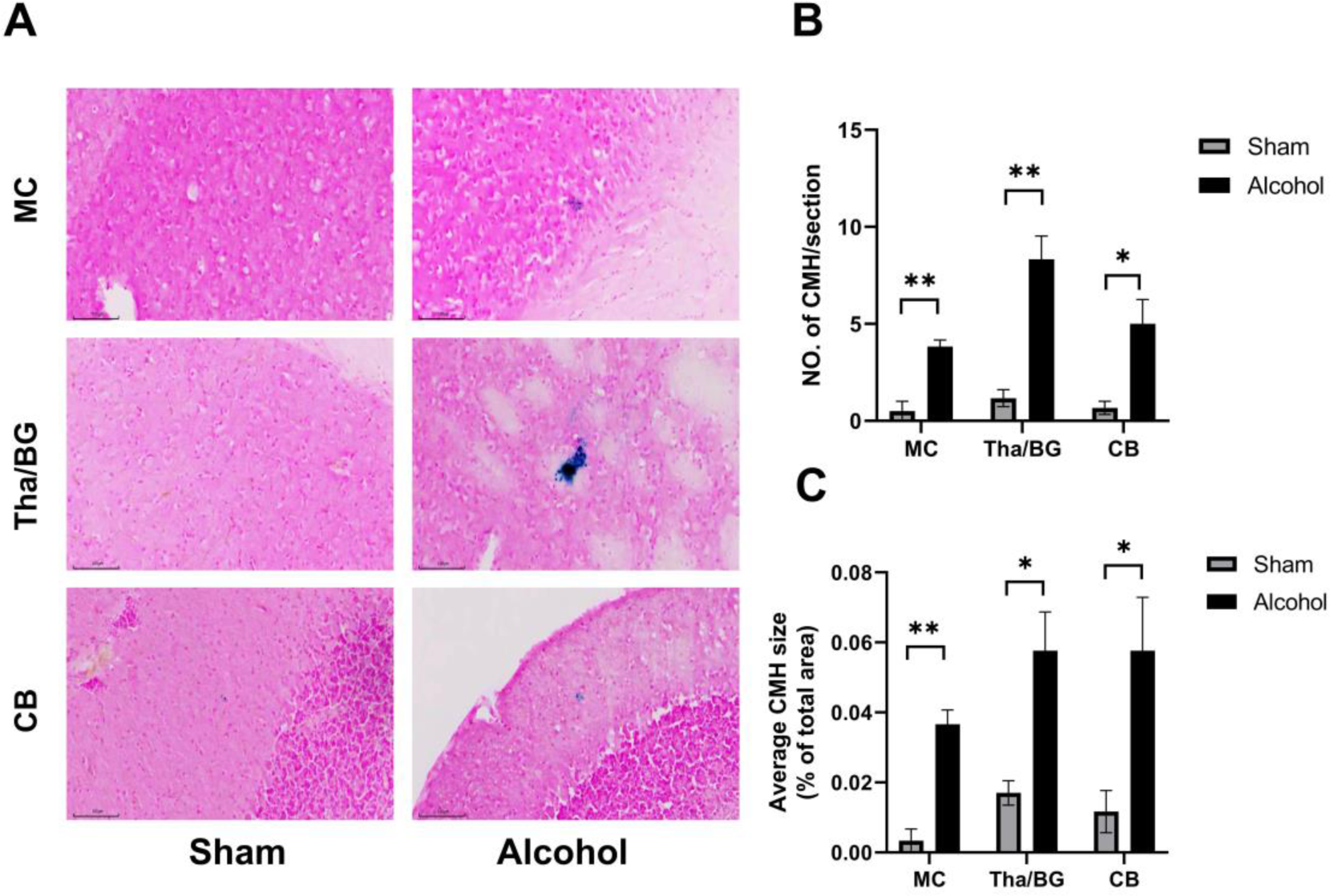
Chronic alcohol exposure induces Prussian Blue-positive cerebral microbleeds. (A) Schematic illustration of Prussian blue staining in motor cortex, thalamus/basal ganglia, and cerebellar tissues. Scale bar=100μm, MC=Motor Cortex, Tha/BG=Thalamus/Basal Ganglia, CB=Cerebellum. (B) Chronic alcohol exposure induces a higher number of cerebral microbleeds. (C) Chronic alcohol exposure induces cerebral microbleeds with larger lesion areas. The data are presented as mean ±S.E.M. (n=3), *p < 0.05, ** p < 0.01, independent samples unpaired t-test.

### 3.2. Chronic alcohol exposure induced motor function decline

After the chronic alcohol exposure model was established, the motor and balance abilities of mice were assessed using the rotarod test. The results showed that after adapting to the equipment, the alcohol group mice had a shorter duration of movement on the rotarod and fell onto the platform more quickly than the control group (Figure.2B), possibly due to poor balance and reduced motor endurance. This suggests that chronic alcohol exposure may lead to a decline in motor function, which could be caused by CMBs.

**Figure.2.**
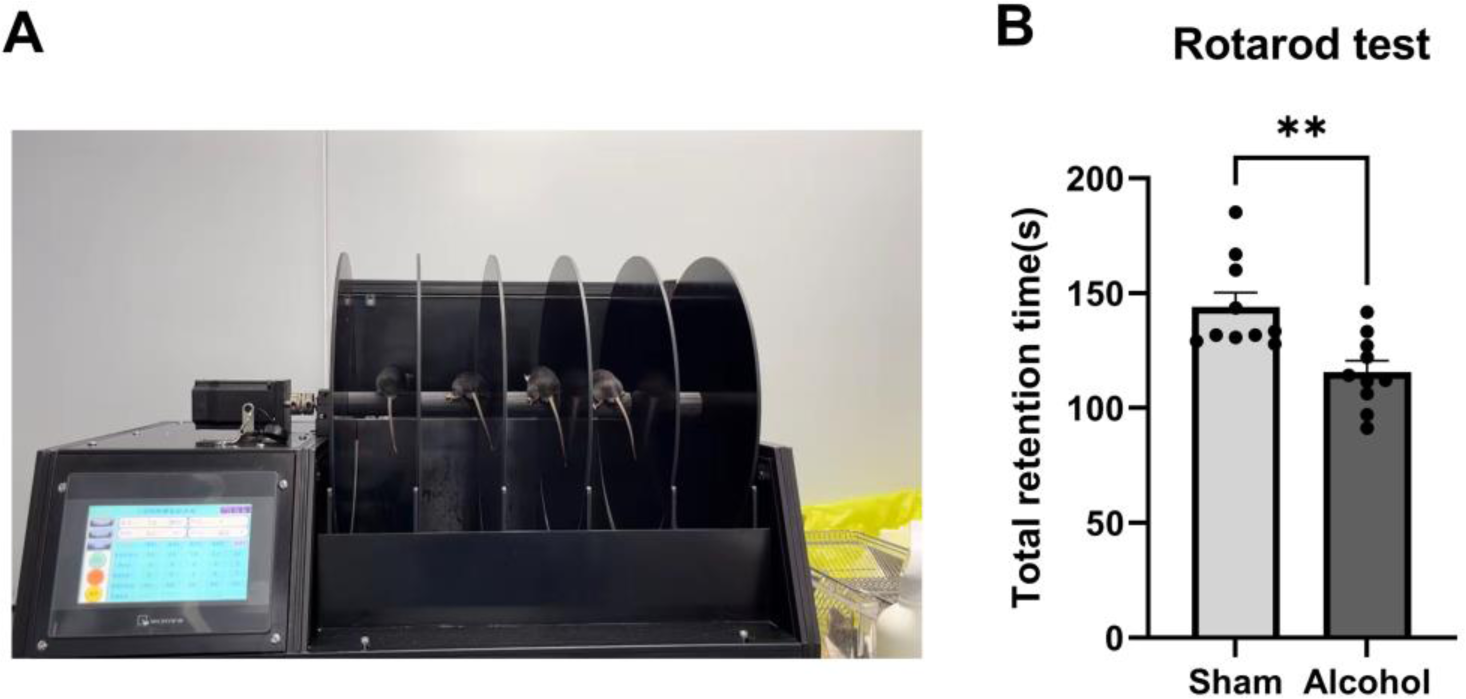
Mice exposed to chronic alcohol show poorer performance in the rotarod test. (A) Schematic illustration of the apparatus used for the rotarod test in this experiment. (B) Total duration of movement on the rotarod. The data are presented as mean ±S.E.M.(n=10), **p=0.0023, independent samples unpaired t-test.

### 3.3. Pirfenidone reduced the excessive deposition of ECM proteins in cerebral small vessels induced by chronic alcohol exposure

Based on the above information, chronic alcohol exposure is known to cause CMBs. We hypothesize that vascular remodeling in cerebral small vessels plays a significant role. Therefore, we used tissue immunofluorescence to co-localize α-SMA, which is specifically expressed in vascular smooth muscle cells, with ECM proteins (collagen I, collagen IV, fibronectin and laminin) deposited on the walls of small vessels. We also added PFD on the basis of chronic alcohol exposure to conduct interventional research. We observed the staining of small vessels in the area shown in Figure.3A. As shown in the Figure.3B, collagen I, collagen IV, fibronectin and laminin are expressed on the walls of small blood vessels, and there are significant differences between the three groups (Sham, Alcohol and Alcohol + PFD). Then we use fluorescence intensity to quantify the expression level of these four proteins (Figure.3C-3F). We found that the deposition of ECM proteins on the cerebral small vessels in the alcohol group was significantly increased compared to the sham group, and intervention group (Alcohol + PFD) reduced the deposition of these proteins.

**Figure.3.**
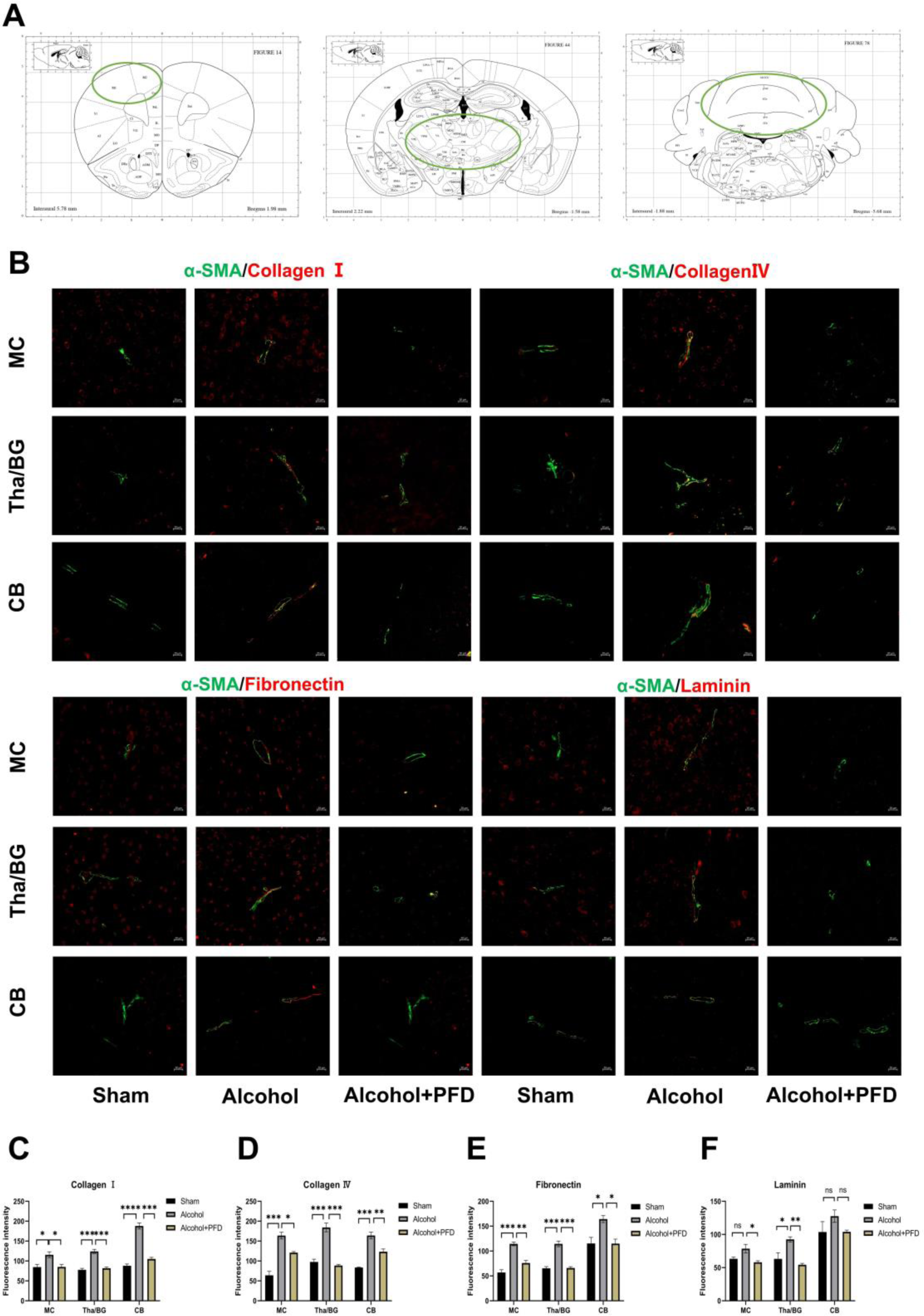
Four types of ECM proteins in the cerebral small vessel in the motor-related regions excessively deposit followed by chronic alcohol exposure. (A) Schematic diagrams of the motor-related brain regions delineated according to the George Paxinos ‘s atlas of mouse brain anatomy. (B) Immunofluorescence staining of four types of ECM proteins: CollagenⅠ(red), CollagenⅣ(red), Fibronectin (red), Laminin (red) co-localized with α-SMA (green) in the MC, Tha/BG, and CB. α-SMA is used to locate cerebral small blood vessels. α-SMA= smooth muscleactin alpha, MC=Motor Cortex, Tha/BG=Thalamus/Basal Ganglia, CB=Cerebellum. Scale bar=20μm. (C) Fluorescence quantitative analysis of CollagenⅠ(MC: Alcohol vs Sham adjust p=0.0435; Alcohol vs Alcohol+PFD adjust p=0.0461; Tha/BG: Alcohol vs Sham adjust p=0.0005; Alcohol vs Alcohol+PFD adjust p=0.0009; CB: Alcohol vs Sham adjust p<0.0001; Alcohol vs Alcohol+PFD adjust p=0.0001. one-way ANOVA with Tukey’s post hoc test), **p=0.002, ***p=0.0003, independent samples unpaired t-test). (D) Fluorescence quantitative analysis of CollagenⅣ(MC: Alcohol vs Sham adjust p=0.0002; Alcohol vs Alcohol+PFD adjust p=0.0178; Tha/BG: Alcohol vs Sham adjust p=0.0005; Alcohol vs Alcohol+PFD adjust p=0.0003; CB: Alcohol vs Sham adjust p=0.0003; Alcohol vs Alcohol+PFD adjust p=0.0087. one-way ANOVA with Tukey’s post hoc test). (E) Fluorescence quantitative analysis of Fibronectin (MC: Alcohol vs Sham adjust p=0.0004; Alcohol vs Alcohol+PFD adjust p=0.0034; Tha/BG: Alcohol vs Sham adjust p=0.0004; Alcohol vs Alcohol+PFD adjust p=0.0004; CB: Alcohol vs Sham adjust p=0.0269; Alcohol vs Alcohol+PFD adjust p=0.0261. one-way ANOVA with Tukey’s post hoc test). (F) Fluorescence quantitative analysis of Laminin (MC: Alcohol vs Sham adjust p=0.0934; Alcohol vs Alcohol+PFD adjust p=0.0316; Tha/BG: Alcohol vs Sham adjust p=0.0280; Alcohol vs Alcohol+PFD adjust p=0.0085; CB: Alcohol vs Sham adjust p=0.3200; Alcohol vs Alcohol+PFD adjust p=0.3249. one-way ANOVA with Tukey’s post hoc test). The data are presented as mean ±SEM (n=3). *p < 0.05, **p < 0.01, ***p < 0.001, ****p < 0.0001. ns non-significant.

### 3.4. Pirfenidone reduced CMBs and improved the motor and balance functions of the mice

At the same time, we used Prussian blue staining (Figure.4A) again to explore whether the use of PFD can reduce the occurrence of microbleeds. As shown in the Figure.4B/4C, the number and size of microbleeds in the brain tissue of mice in the intervention group (alcohol + PFD) were reduced compared to the alcohol group. Furthermore, we used SWI sequence of small animal magnetic resonance to detect microbleeds. As shown in the Figure.4E, circular-like low signal lesions appeared in the deep brain of mice exposed to chronic alcohol, which were not found in the control group and the intervention group. Rotarod tests demonstrated that mice orally administered with PFD had a longer duration of movement on the rotarod than those in the alcohol-only group, suggesting an improvement in balance and motor functions.

**Figure.4.**
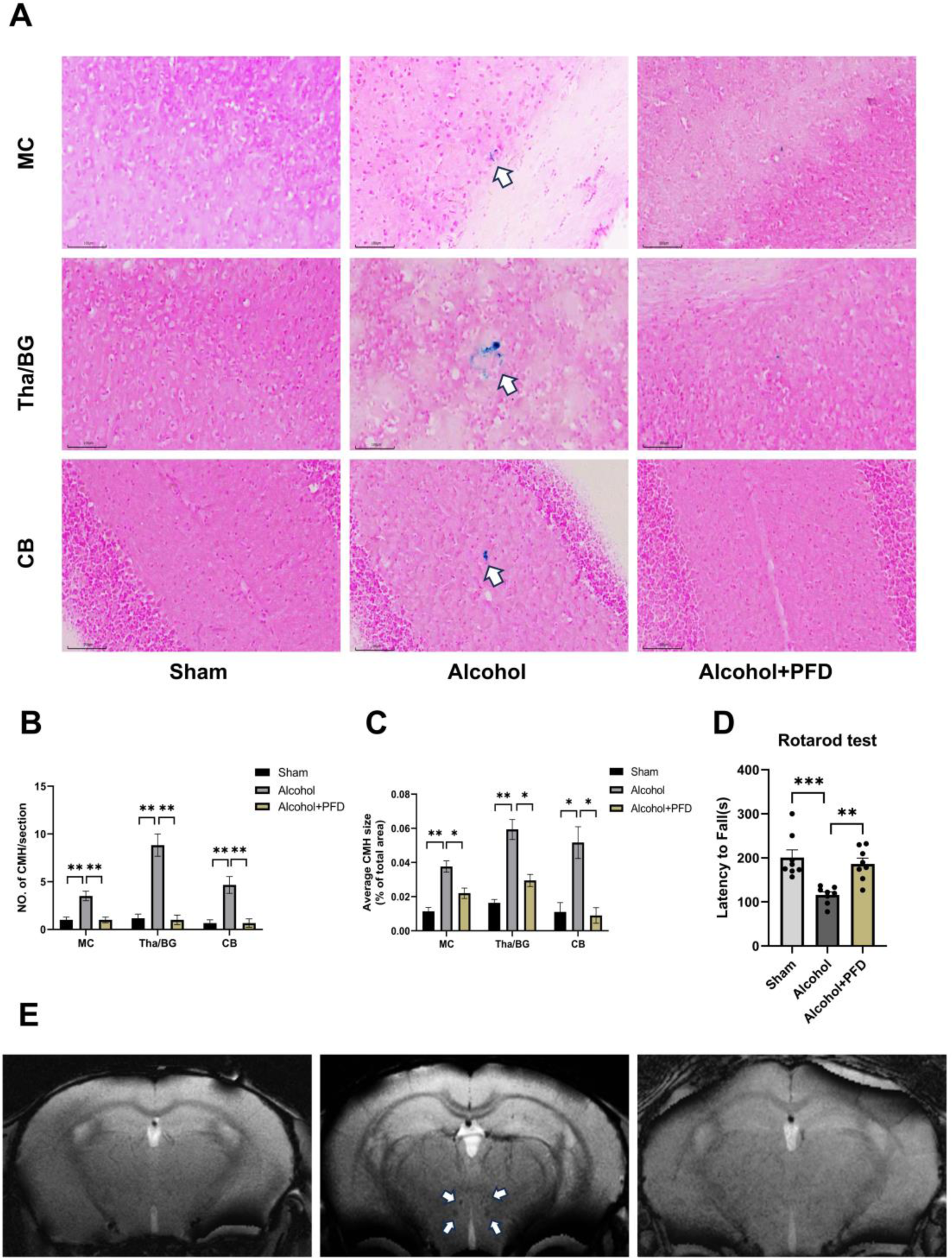
The improvement in microhemorrhage and motor function in mice after PFD intervention. (A) Schematic illustration of Prussian blue staining in motor cortex, thalamus/basal ganglia, and cerebellar of three groups (Sham, Alcohol and Alcohol + PFD). Scale bar=100μm, MC=Motor Cortex, Tha/BG=Thalamus/Basal Ganglia, CB=Cerebellum, PFD=Pirfenidone. (B) Average of CMHs size in different regions of groups. (MC: Alcohol vs Sham adjust p=0.0017; Alcohol vs Alcohol +PFD adjust p=0.0203; Tha/BG: Alcohol vs Sham adjust p=0.0016; Alcohol vs Alcohol +PFD adjust p=0.0133; CB: Alcohol vs Sham adjust p=0.0129; Alcohol vs Alcohol +PFD adjust p=0.0103. one-way ANOVA with Tukey’s post hoc test). (C) The number of CMHs in different regions of groups. (MC: Alcohol vs Sham adjust p=0.0076; Alcohol vs Alcohol +PFD adjust p=0.0076; Tha/BG: Alcohol vs Sham adjust p=0.0026; Alcohol vs Alcohol +PFD adjust p=0.0039; CB: Alcohol vs Sham adjust p=0.0079; Alcohol vs Alcohol +PFD adjust p=0.0079. one-way ANOVA with Tukey’s post hoc test). (D) Total duration of movement on the rotarod. (Alcohol vs Sham adjust p=0.0005; Alcohol vs Alcohol +PFD adjust p=0.0028. one-way ANOVA with Tukey’s post hoc test). (E) Small animal magnetic resonance SWI sequence imaging. The white arrow points to the location of signal abnormalities. The data are presented as mean ±SEM. *p < 0.05, **p < 0.01, ***p < 0.001, ****p < 0.0001. ns non-significant.

### 3.5. Pirfenidone administration alleviated alcohol-induced phenotypic transformation of cerebral microvascular smooth muscle cells

Next, we delved further into the reasons behind the abnormal accumulation of ECM proteins on the walls of brain small vessels and induced by alcohol. We know that the ECM is primarily synthesized by vascular smooth muscle cells. Therefore, we treated CMVSMCs with gradient concentrations of alcohol and then tested cell viability using the CCK-8 assay kit. The results showed that alcohol inhibited CMVSMCs in a concentration- and time-dependent manner. Ultimately, we chose a treatment scheme of 100mM ethanol solution for 24 hours. Finally, the CMVSMCs were divided into three groups: control, alcohol (100mM, ethanol, 24 hours), and intervention group (100mM, ethanol, 24 hours; 0.3mg/ml, Pirfenidone, 24 hours). After treatment, we found that the long diameter of CMVSMCs treated with alcohol was shortened, which prolonged upon adding Pirfenidone (Figure.5A). We converted the microscopic images into binary images to more intuitively reflect the changes in cell morphology (Figure.5B). Subsequently, qPCR and WB results showed that the expression of Decorin and Lumican in CMVSMCs treated with alcohol was upregulated, consistent with the characteristics of the transformation of smooth muscle cells into fibroblast-like phenotypes. Moreover, the expression of these two markers was downregulated upon adding Pirfenidone (Figure.5C-5G).

**Figure.5.**
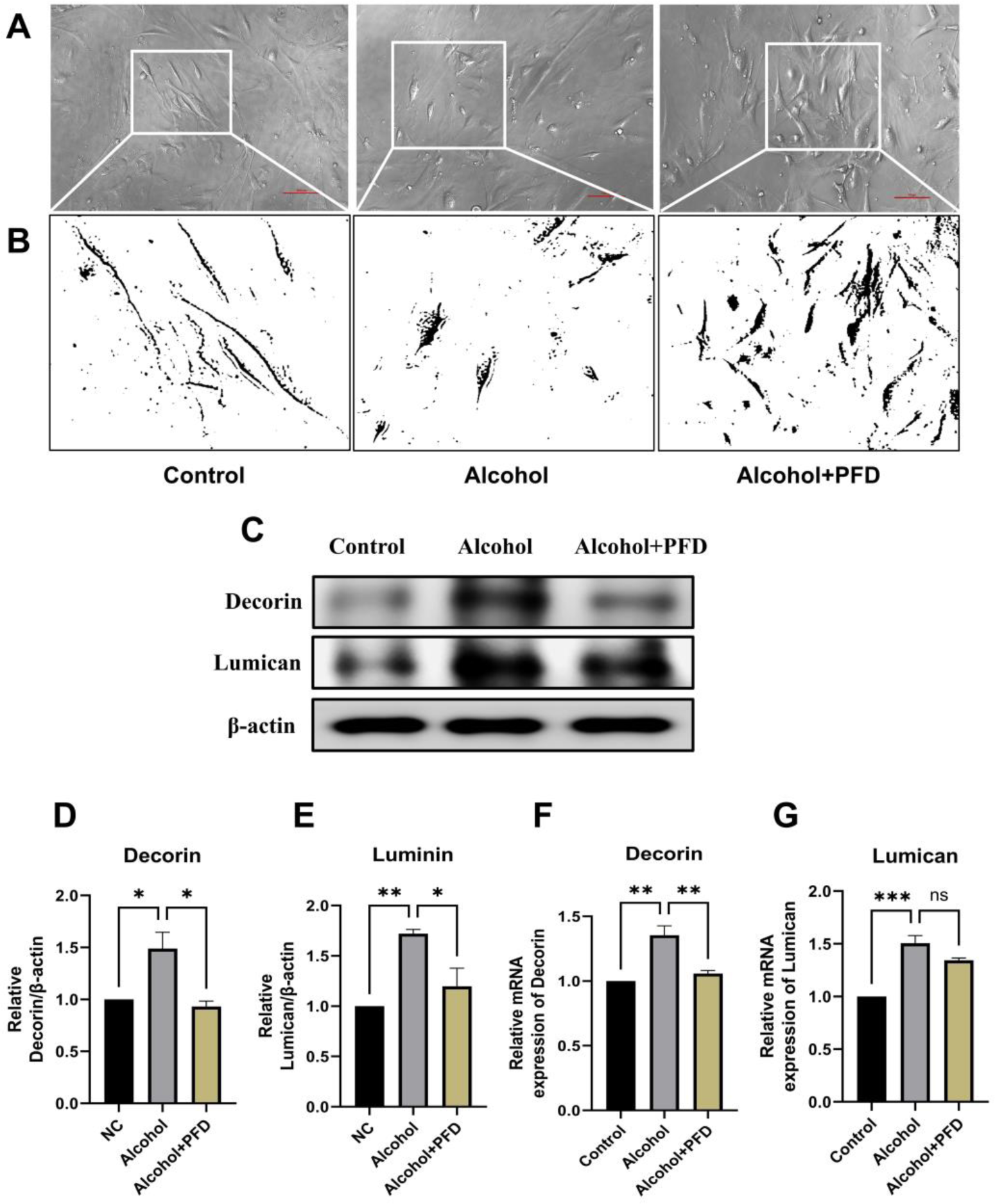
Changes in morphology and phenotypic transformation markers in CMVSMCs after alcohol and PFD treatment. (A) CMVSMCs morphology after treatment of alcohol or PFD. The white box indicates the area shown in Figure.5b. (B) The binary image within the white box area in Figure.5a. (C) Western blotting analysis of Decorin and Lumican. (D) Western blot of Decorin (Alcohol vs Control adjust p=0.0251; Alcohol vs Alcohol +PFD adjust p=0.0142, one-way ANOVA with Tukey’s post hoc test). (E) Western blot of Lumican (Alcohol vs Control adjust p=0.0071; Alcohol vs Alcohol +PFD adjust p=0.0302, one-way ANOVA with Tukey’s post hoc test) (F) qPCR analysis of Decorin (Alcohol vs Control adjust p=0.0032; Alcohol vs Alcohol +PFD adjust p=0.0078, one-way ANOVA with Tukey’s post hoc test). (G) qPCR analysis of Lumican (Alcohol vs Control adjust p=0.0004; Alcohol vs Alcohol +PFD adjust p=0.0817, one-way ANOVA with Tukey’s post hoc test). The data are presented as mean ±SEM. *p < 0.05, **p < 0.01, ***p < 0.001, ****p < 0.0001. ns non-significant.

### 3.6. Pirfenidone inhibited the alcohol-induced overproduction of ECM proteins in CMVSMCs by suppressing TGFβ1/Smad signaling pathway

Then, we pondered whether this phenotypic transformation of the cells is also accompanied by changes in their synthesis function. Therefore, through cell immunofluorescence, qPCR, and WB, we examined the expression of ECM proteins in CMVSMCs. The results (Figure.6A-6N) showed that the synthesis of ECM proteins in CMVSMCs increased after alcohol treatment, which is consistent with the results of animal experiments. Previous studies have shown that after the activation of the TGFβ1/Smad pathway, it stimulates the synthesis of downstream collagen and fibronectin. Therefore, we also examined the expression of TGFβ1, Smad, and pSmad in CMVSMCs. The results (Figure.6O-6U) showed that the expression of these proteins in CMVSMCs of the alcohol group was upregulated, indicating that the signaling pathway was activated. And after the addition of Pirfenidone, the pathway was inhibited, and the expression of ECM proteins was correspondingly reduced. This suggests that CMVSMCs might be involved in the abnormal deposition of ECM proteins on the vascular walls after alcohol treatment, mediated by the TGFβ1/Smad signaling pathway. Pirfenidone, as a specific inhibitor of this signaling pathway, can reverse the aforementioned process.

**Figure.6.**
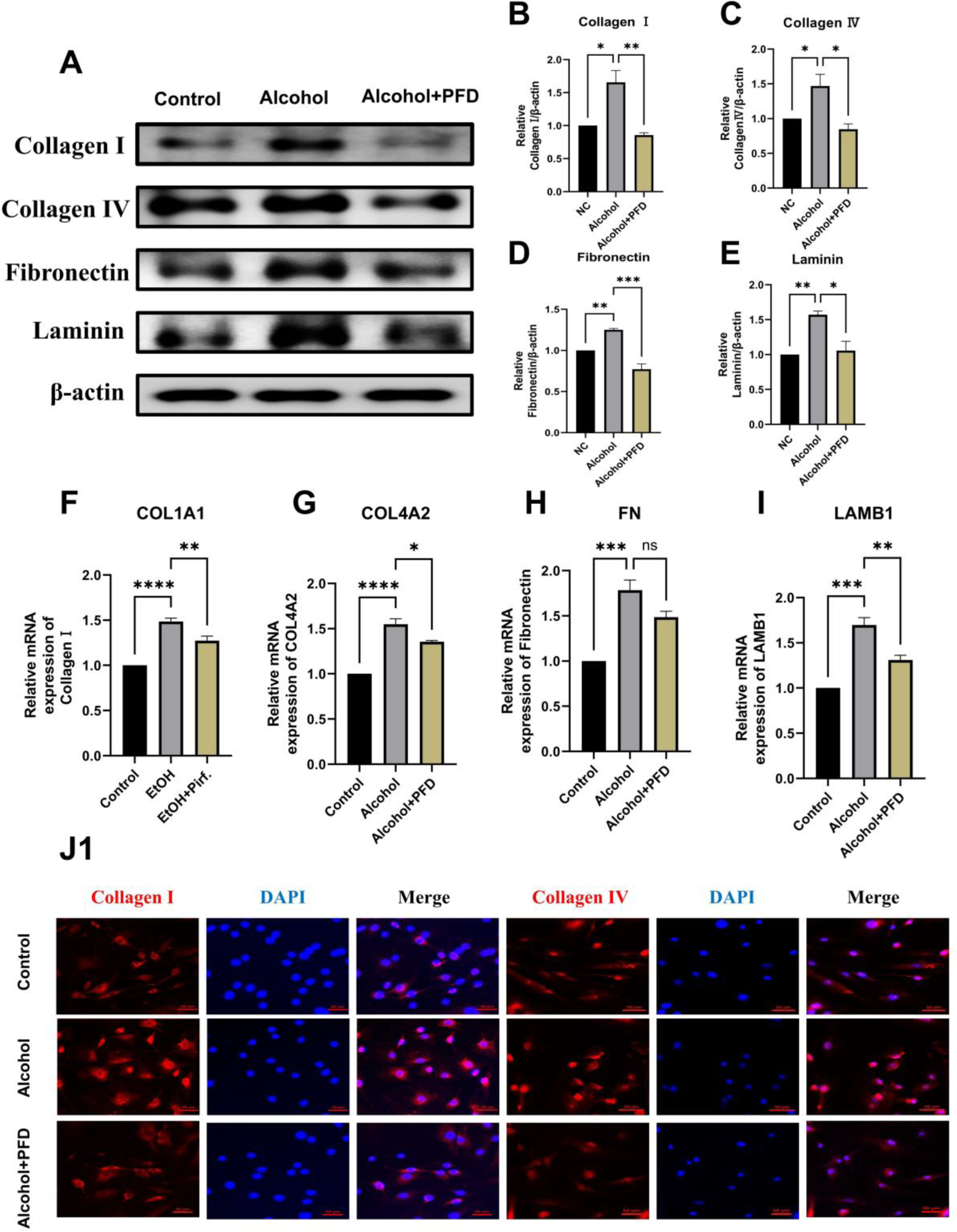
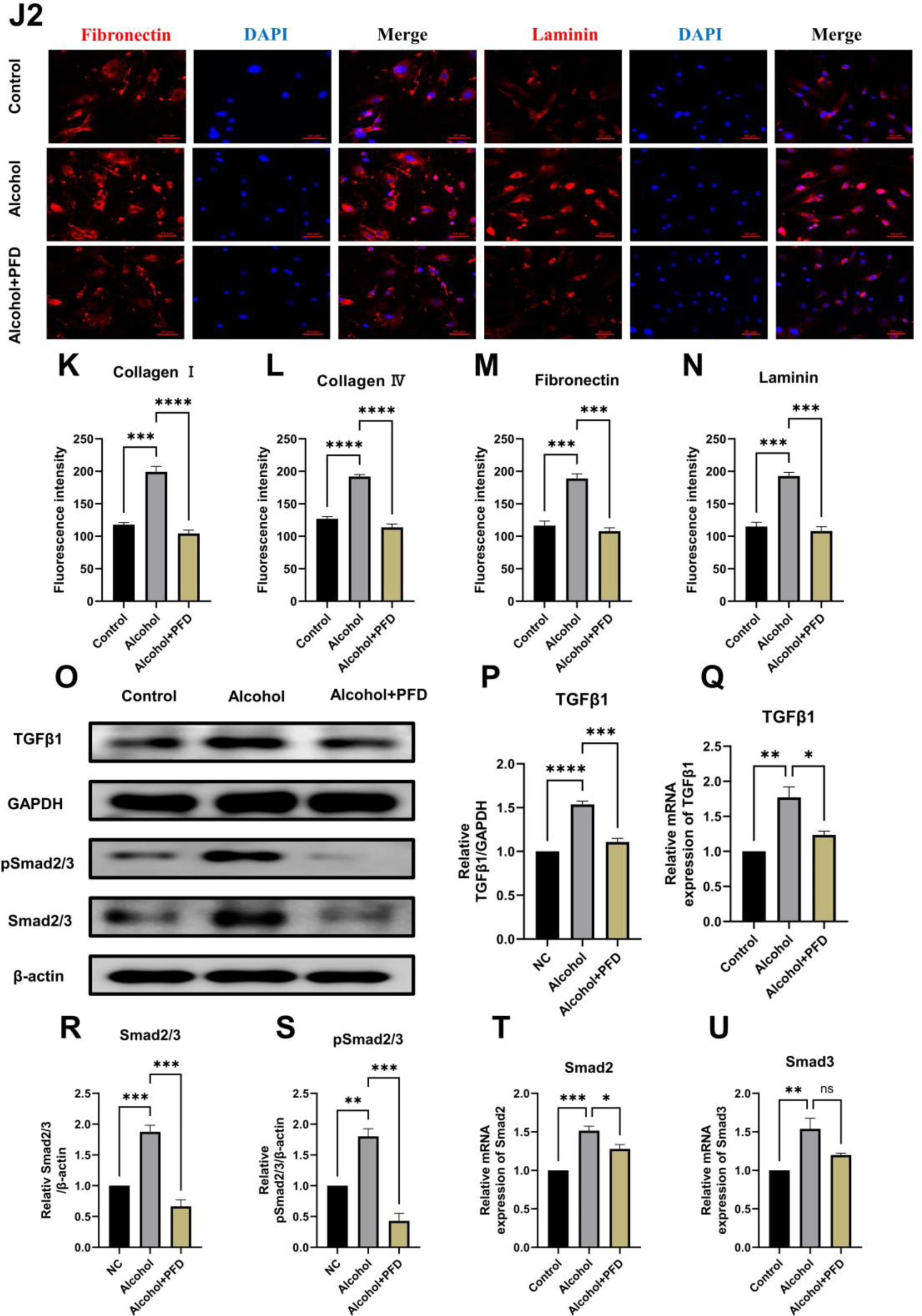
Expression of ECMs and TGFβ1/Smad of CMVSMCs after the treatment of alcohol or PFD. (A) Western blotting analysis of CollagenⅠ, CollagenⅣ, Fibronectin and Laminin. (B) Western blot of CollagenⅠ (Alcohol vs Control adjust p=0.0107; Alcohol vs Alcohol +PFD adjust p=0.0041, one-way ANOVA with Tukey’s post hoc test). (C) Western blot of CollagenⅣ (Alcohol vs Control adjust p=0.0485; Alcohol vs Alcohol +PFD adjust p=0.0149, one-way ANOVA with Tukey’s post hoc test). (D) Western blot of Fibronectin (Alcohol vs Control adjust p=0.0094; Alcohol vs Alcohol +PFD adjust p=0.0003, one-way ANOVA with Tukey’s post hoc test). (E) Western blot of Laminin (Alcohol vs Control adjust p=0.0066; Alcohol vs Alcohol +PFD adjust p=0.0110, one-way ANOVA with Tukey’s post hoc test). (F) qPCR analysis of CollagenⅠ (Alcohol vs Control adjust p<0.0001; Alcohol vs Alcohol +PFD adjust p=0.0059, one-way ANOVA with Tukey’s post hoc test). (G) qPCR analysis of CollagenⅣ (Alcohol vs Control adjust p<0.0001; Alcohol vs Alcohol +PFD adjust p=0.0214, one-way ANOVA with Tukey’s post hoc test). (H) qPCR analysis of Fibronectin (Alcohol vs Control adjust p=0.0007; Alcohol vs Alcohol +PFD adjust p=0.0671, one-way ANOVA with Tukey’s post hoc test). (I) qPCR analysis of Laminin (Alcohol vs Control adjust p=0.0003; Alcohol vs Alcohol +PFD adjust p=0.0061, one-way ANOVA with Tukey’s post hoc test). (J) Immunofluorescence staining of four types of ECM proteins: CollagenⅠ(red), CollagenⅣ(red), Fibronectin (red), Laminin (red) and DAPI (blue). Bar scale=50μm. (K) Fluorescence quantitative analysis of CollagenⅠ(Alcohol vs Control adjust p=0.0002; Alcohol vs Alcohol +PFD adjust p<0.0001, one-way ANOVA with Tukey’s post hoc test). (L) Fluorescence quantitative analysis of CollagenⅣ (Alcohol vs Control adjust p<0.0001; Alcohol vs Alcohol +PFD adjust p<0.0001, one-way ANOVA with Tukey’s post hoc test). (M) Fluorescence quantitative analysis of Fibronectin (Alcohol vs Control adjust p=0.0005; Alcohol vs Alcohol +PFD adjust p=0.0003, one-way ANOVA with Tukey’s post hoc test). (N) Fluorescence quantitative analysis of Laminin (Alcohol vs Control adjust p=0.0003; Alcohol vs Alcohol +PFD adjust p=0.0002, one-way ANOVA with Tukey’s post hoc test). (O) Western blotting analysis of TGFβ1, Smad2/3 and phosphor-Smad2/3. (P) Western blot of TGFβ1 (Alcohol vs Control adjust p<0.0001; Alcohol vs Alcohol +PFD adjust p=0.0002, one-way ANOVA with Tukey’s post hoc test). (Q) qPCR analysis of TGFβ1 (Alcohol vs Control adjust p=0.0024; Alcohol vs Alcohol +PFD adjust p=0.0143, one-way ANOVA with Tukey’s post hoc test). (R) Western blot of Smad2/3 (Alcohol vs Control adjust p=0.0009; Alcohol vs Alcohol +PFD adjust p=0.0001, one-way ANOVA with Tukey’s post hoc test). (S) Western blot of pSmad2/3 (Alcohol vs Control adjust p=0.0030; Alcohol vs Alcohol +PFD adjust p=0.0002, one-way ANOVA with Tukey’s post hoc test). (T) qPCR analysis of Smad2 (Alcohol vs Control adjust p=0.0006; Alcohol vs Alcohol +PFD adjust p=0.0273, one-way ANOVA with Tukey’s post hoc test). (U) qPCR analysis of Smad3 (Alcohol vs Control adjust p=0.0076; Alcohol vs Alcohol +PFD adjust p=0.0551, one-way ANOVA with Tukey’s post hoc test). The data are presented as mean ±SEM. *p < 0.05, **p < 0.01, ***p < 0.001, ****p < 0.0001. ns non-significant.

**Figure.7.**
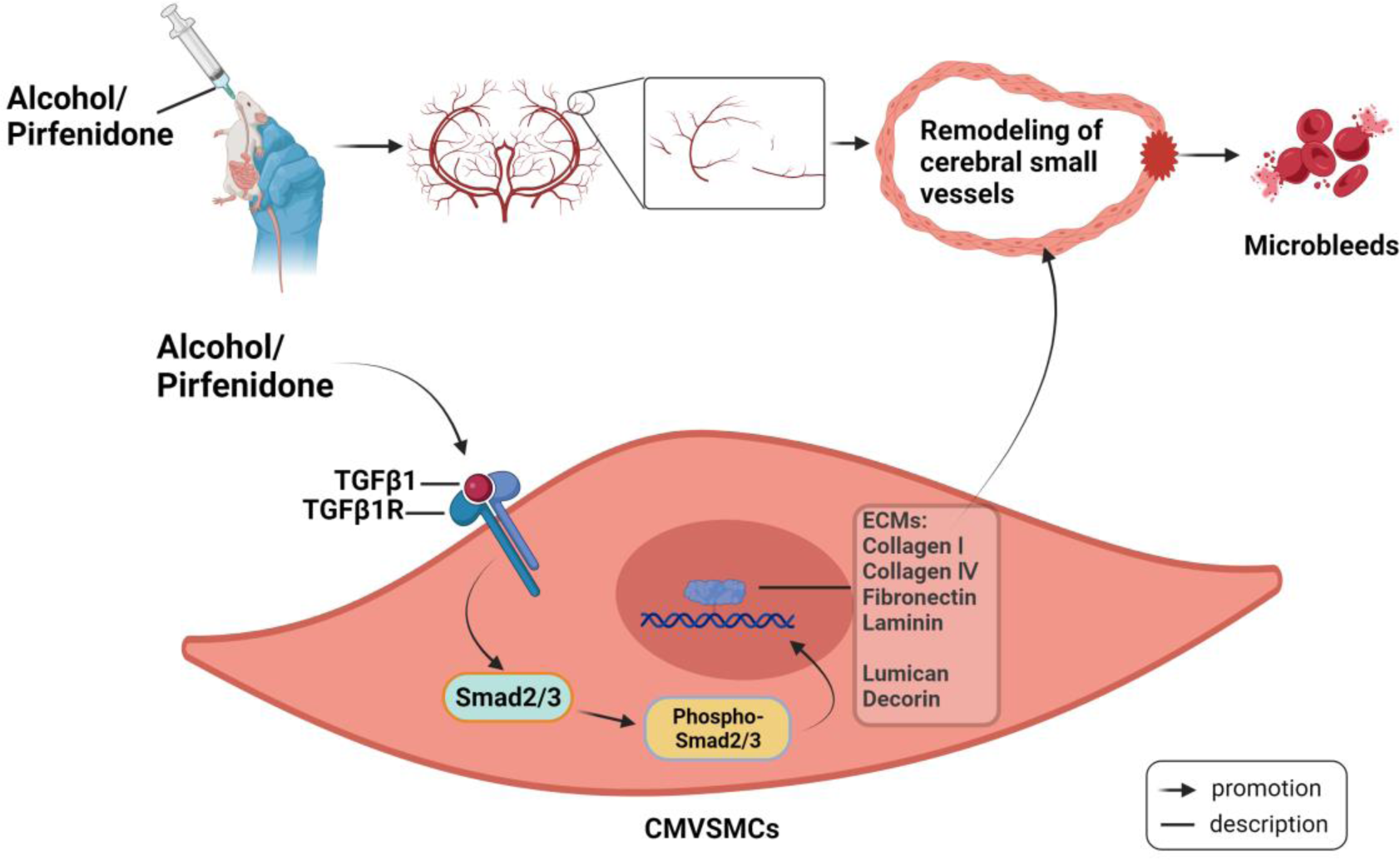
A probable overview of alcohol-induced cerebral microbleeds and its underlying mechanisms. Alcohol can indeed directly induce cerebral microbleeds in mice, accompanying by cerebral microvascular remodeling mediated by the TGF-β1/Smad pathway in cerebral microvascular smooth muscle cells. Abnormal deposition of ECM leads to increased vascular stiffness and decreased compliance, resulting in vascular rupture and bleeding. These may explain the formation of cerebral microbleeds.

## 4. Discussion

CMBs are a radiological manifestation of CSVD^4^. There have been many community epidemiological studies exploring the prevalence and high-risk factors of CMBs, as well as clinical studies based on specific clinical diseases, such as cerebral amyloid angiopathy^45^ and Alzheimer’s disease^46^. Among them, there are some large-scale clinical studies on the relationship between alcohol consumption and CMBs. From those studies, it can be concluded that alcohol consumption, especially heavy drinking, increases the risk of CMBs, and is more strongly associated with deep CMBs^6,7,9^. However, there are still relatively few studies that explore the mechanisms underlying the formation of microbleeds. The transition from radiological findings to in-depth histopathological studies remains a challenge. Our study preliminarily found that in the chronic alcohol consumption mouse model we constructed, histological and radiological evidence of CMBs was found using Prussian blue staining and small animal magnetic resonance scanning. However, the bleeding points are relatively scattered, and the responsible small blood vessels have not yet been identified. This may be related to the insufficient restoration of the spatial structure in the coronal sections.

CSVD is believed to be associated with gait abnormalities, particularly in cases of white matter hyperintensities and lacunar infarcts^47,48^. Clinical studies have also reported an association between CMBs and impaired gait and balance^27,28^, but there is a relative lack of basic research data in this area. Therefore, after discovering that alcohol can induce CMBs, we focused our behavioral and molecular biology researches on motor functions and brain regions related to movement. In addition to the frontal lobes and deep brain regions (such as the thalamus and basal ganglia) involved in the studies mentioned above, we also included the cerebellum in our research scope, as it is a key area controlling body balance and coordination. Considering the frontal lobe’s involvement in numerous neural circuits, we referred to George Paxinos ‘s atlas of mouse brain anatomy, focusing on the motor cortex of the frontal lobe (primary and secondary motor cortex) which is more closely related to motor functions. For behavioral testing, we chose one of the most commonly used kinematic experiments, the rotarod test^38^, to measure the mice’s motor and balance abilities. Our results showed that alcohol-exposed mice had microbleeds in the tissues of the MC, Tha/BG, and CB, higher than in the control group. Moreover, mice exposed to chronic alcohol performed poorly in the rotarod test, falling off the platform more quickly. Considering that the handling and gavage during the administration could potentially injure the mice’s limbs and joints, our control group mice were subjected to the same handling and gavage procedures with water, thus our results account for potential biases due to manual operations and are credible.

Small vessel remodeling has long been considered one of the pathogenic mechanisms of CSVD, with arteriolosclerosis being the most common pathological subtype^12^. CMBs, a type of CSVD, are closely associated with arteriolosclerosis^49^. Liu, N., et al. observed in a hypertension-induced CSVD model that small vessels exhibited excessive deposition of collagen, fibronectin, and laminin, leading to narrowing of the lumen and reduced cerebral perfusion^13^. Therefore, in our experiment, we observed the excessive deposition of ECM proteins in the cerebral microvessels of mice (MC, Tha/BG, and CB). This abnormal deposition contributes to the progression of arteriolosclerosis.

Cerebral small blood vessels are the primary resistance vessels that distribute blood flow to the capillary bed, where the greatest changes in blood pressure and flow velocity occur. This is crucial for maintaining adequate blood flow in the subcortical structures of the brain^50,51^. Recent studies have shown that the control of cerebral blood flow is primarily due to the action of vascular smooth muscle cells, rather than pericytes, which aligns with previous understandings^52^. The distribution of cerebral vascular smooth muscle cells terminates at the cerebral small arteries and arterioles, where there are typically one to two layers of smooth muscle cells^53^. These cells mainly perform two functions: first, they maintain vascular tension, regulating the contraction and relaxation of blood vessels, thereby controlling cerebral blood flow; second, they synthesize and renew ECM proteins like collagen and elastin, ensuring the structural integrity of the vascular wall^54^. Previous research has confirmed that alcohol can affect the state of vascular contraction and relaxation by altering the voltage-gated calcium channels, BK channels, and magnesium channels in cerebral vascular smooth muscle cells, thereby influencing the electrolyte balance inside and outside the cells and ultimately affecting cerebral perfusion^55–57^. Therefore, our study mainly focuses on the second function, namely, the synthesis of ECM proteins. We wondered if alcohol also stimulates CMVSMCs to synthesize excessive ECM proteins, leading to their deposition in the vascular wall, and our preliminary results have confirmed this. We also noted that smooth muscle cells are often affected by inflammation, intercellular interactions, mechanical stress, growth factors, and transforming factors, leading to phenotypic transformation^19^. Literature review indicates that the change in the synthetic function of CMVSMCs is a result of their phenotypic transformation^20^. Hence, we isolated primary CMVSMCs and indeed found evidence of phenotypic transformation and the induced synthesis of excessive ECM due to this transformation.

TGFβ signaling promotes the differentiation, maturation, proliferation, migration, and adhesion of smooth muscle cells^58^. Under pathological conditions, both astrocytes and microglia can secrete excessive TGFβ, and it is speculated that this may be directly related to vascular damage^59^. Our research has found that the expression of TGFβ1 is also upregulated in vascular smooth muscle cells under the influence of alcohol. In the cerebral vessels of TGFβ1 transgenic mice, an increase in basement membrane proteins, perlecan, and fibronectin was observed^60^, suggesting that TGFβ1 signaling may be related to the synthesis of ECM proteins. Our experiments also show that the upregulated TGFβ1 can activate downstream Smad2/3. The activated, phosphorylated Smad2/3 enters the nucleus and then promotes the initiation of the synthesis of ECM proteins.

Additionally, these vascular remodeling could lead to impaired clearance of toxic substances, resulting in the deposition of amyloid proteins in the vessel walls. It is well known that amyloid proteins are damaging to vascular cells and structures^61^. Thus, the activation of TGFβ1 signaling induced by alcohol may have broader implications. Whether other cell types, such as microglia and astrocytes, are also involved in this process and the underlying mechanisms, remains to be further explored.

In this study, we also conducted interventional research using a novel drug, pirfenidone, which specifically inhibits the TGFβ1/Smad pathway. Pirfenidone has been used in the central nervous system, including in traumatic brain injuries, where it acts as a neuroprotectant and anti-inflammatory agent, reducing mortality and morbidity post-injury^62^. It’s also a preventive medication for secondary hemorrhage and cerebral infarction after subarachnoid hemorrhage^63^. Applying it to our animal model, we observed a reduction in ECM protein deposition in small vessels and improvements in microbleeding and motor balance functions upon successful inhibition of the TGFβ1/Smad pathway. We administered pirfenidone via gavage, a common method in animal studies, as it can cross the blood-brain barrier. However, for central system disease research, stereotactic in situ injection might be a more direct method of administration.

## 5. Conclusion

In conclusion, our study demonstrates that chronic alcohol exposure can induce microbleeds and impair motor balance abilities. This is associated with vascular remodeling caused by excessive deposition of ECM proteins in the walls of small blood vessels. The activation of the TGFβ1/Smad signaling pathway in vascular smooth muscle cells mediates this process, and pirfenidone shows promising therapeutic effects. Therefore, TGFβ1 may be a potential therapeutic target for alcohol-related CMBs or small vessel disease.

## Acknowledgements

This work was supported by the Grant of China and Germany Collaboration & Exchange Project (NO: M0746) and partly supported by the Grant of National Key R&D Program of China (2018YFC1314400, 2018YFC1314401 to Y.P)

## References

1. Hendriks HFJ. Alcohol and Human Health: What Is the Evidence? Annu Rev Food Sci Technol. 2020;11:1–21

2. Global status report on alcohol and health 2018. 2023(2023-12-17):This report presents a comprehensive picture of alcohol consumption, disease burden and policy responses worldwide.

3. Millwood IY, Walters RG, Mei XW et al. Conventional and genetic evidence on alcohol and vascular disease aetiology: a prospective study of 500 000 men and women in China. Lancet (London, England). 2019;393(10183):1831–1842

4. Greenberg SM, Vernooij MW, Cordonnier C et al. Cerebral microbleeds: a guide to detection and interpretation. The Lancet. Neurology. 2009;8(2):165–74

5. Cheng A, Batool S, McCreary CR et al. Susceptibility-weighted imaging is more reliable than T2*-weighted gradient-recalled echo MRI for detecting microbleeds. Stroke. 2013;44(10):2782–6

6. Haller S, Vernooij MW, Kuijer JPA et al. Cerebral Microbleeds: Imaging and Clinical Significance. Radiology. 2018;287(1):11–28

7. Copenhaver BR, Hsia AW, Merino JG et al. Racial differences in microbleed prevalence in primary intracerebral hemorrhage. Neurology. 2008;71(15):1176–82

8. Lu D, Liu J, MacKinnon AD, Tozer DJ, Markus HS. Prevalence and Risk Factors of Cerebral Microbleeds: An Analysis From the UK Biobank. Neurology. 2021;

9. Ding J, Sigurdsson S, Garcia M et al. Risk Factors Associated With Incident Cerebral Microbleeds According to Location in Older People: The Age, Gene/Environment Susceptibility (AGES)-Reykjavik Study. JAMA Neurol. 2015;72(6):682–8

10. Cannistraro RJ, Badi M, Eidelman BH et al. CNS small vessel disease: A clinical review. Neurology. 2019;92(24):1146–1156

11. Pantoni L. Cerebral small vessel disease: from pathogenesis and clinical characteristics to therapeutic challenges. The Lancet. Neurology. 2010;9(7):689–701

12. Ter Telgte A, van Leijsen EMC, Wiegertjes K et al. Cerebral small vessel disease: from a focal to a global perspective. Nature reviews. Neurology. 2018;14(7):387–398

13. Liu N, Tang J, Xue Y et al. EP3 Receptor Deficiency Improves Vascular Remodeling and Cognitive Impairment in Cerebral Small Vessel Disease. Aging Dis. 2022;13(1):313–328

14. Laurent S, Boutouyrie P. The structural factor of hypertension: large and small artery alterations. Circ Res. 2015;116(6):1007–21

15. Frösen J, Joutel A. Smooth muscle cells of intracranial vessels: from development to disease. Cardiovasc Res. 2018;114(4):501–512

16. Li W, Li J, Liu W, Altura BT, Altura BM. Alcohol-induced apoptosis of canine cerebral vascular smooth muscle cells: role of extracellular and intracellular calcium ions. Neurosci Lett. 2004;354(3):221–4

17. Yang ZW, Wang J, Zheng T, Altura BT, Altura BM. Ethanol-induced contractions in cerebral arteries: role of tyrosine and mitogen-activated protein kinases. Stroke. 2001;32(1):249–57

18. Liu P, Xi Q, Ahmed A, Jaggar JH, Dopico AM. Essential role for smooth muscle BK channels in alcohol-induced cerebrovascular constriction. Proc Natl Acad Sci U S A. 2004;101(52):18217–22

19. Owens GK, Kumar MS, Wamhoff BR. Molecular regulation of vascular smooth muscle cell differentiation in development and disease. Physiol Rev. 2004;84(3):767–801

20. Shi J, Yang Y, Cheng A, Xu G, He F. Metabolism of vascular smooth muscle cells in vascular diseases. American journal of physiology. Heart and circulatory physiology. 2020;319(3):H613–H631

21. Wirka RC, Wagh D, Paik DT et al. Atheroprotective roles of smooth muscle cell phenotypic modulation and the TCF21 disease gene as revealed by single-cell analysis. Nat Med. 2019;25(8):1280–1289

22. Basatemur GL, Jørgensen HF, Clarke MCH, Bennett MR, Mallat Z. Vascular smooth muscle cells in atherosclerosis. Nature reviews. Cardiology. 2019;16(12):727–744

23. Ruiz-Ortega M, Rodríguez-Vita J, Sanchez-Lopez E, Carvajal G, Egido J. TGF-beta signaling in vascular fibrosis. Cardiovasc Res. 2007;74(2):196–206

24. Meng X, Nikolic-Paterson DJ, Lan HY. TGF-β: the master regulator of fibrosis. Nature reviews. Nephrology. 2016;12(6):325–38

25. Ciuclan L, Ehnert S, Ilkavets I et al. TGF-beta enhances alcohol dependent hepatocyte damage via down-regulation of alcohol dehydrogenase I. J Hepatol. 2010;52(3):407–16

26. Chen N, Geng Q, Zheng J et al. Suppression of the TGF-β/Smad signaling pathway and inhibition of hepatic stellate cell proliferation play a role in the hepatoprotective effects of curcumin against alcohol-induced hepatic fibrosis. Int J Mol Med. 2014;34(4):1110–6

27. de Laat KF, van den Berg HAC, van Norden AGW, et al. Microbleeds are independently related to gait disturbances in elderly individuals with cerebral small vessel disease. Stroke. 2011;42(2):494–7

28. Mao H, Zhang J, Zhu W et al. Basal Ganglia and Brainstem Located Cerebral Microbleeds Contributed to Gait Impairment in Patients with Cerebral Small Vessel Disease. Journal of Alzheimer’s disease : JAD. 2023;94(3):1005–1012

29. Mistarz N, Canfield L, Nielsen DG, Skøt L, Mellentin AI. Gait ataxia in alcohol use disorder: A systematic review. Psychology of addictive behaviors : journal of the Society of Psychologists in Addictive Behaviors. 2023;

30. Lan L, Wang H, Zhang X et al. Chronic exposure of alcohol triggers microglia-mediated synaptic elimination inducing cognitive impairment. Exp Neurol. 2022;353:114061

31. Liu D, Li J, Rong X et al. Cdk5 Promotes Mitochondrial Fission via Drp1 Phosphorylation at S616 in Chronic Ethanol Exposure-Induced Cognitive Impairment. Mol Neurobiol. 2022;59(12):7075–7094

32. Carson EJ, Pruett SB. Development and characterization of a binge drinking model in mice for evaluation of the immunological effects of ethanol. Alcoholism, clinical and experimental research. 1996;20(1):132–8

33. Shihab FS, Bennett WM, Yi H, Andoh TF. Pirfenidone treatment decreases transforming growth factor-beta1 and matrix proteins and ameliorates fibrosis in chronic cyclosporine nephrotoxicity. American journal of transplantation : official journal of the American Society of Transplantation and the American Society of Transplant Surgeons. 2002;2(2):111–9

34. Kakugawa T, Mukae H, Hayashi T et al. Pirfenidone attenuates expression of HSP47 in murine bleomycin-induced pulmonary fibrosis. The European respiratory journal. 2004;24(1):57–65

35. DUNHAM NW, MIYA TS. A note on a simple apparatus for detecting neurological deficit in rats and mice. Journal of the American Pharmaceutical Association. American Pharmaceutical Association. 1957;46(3):208–9

36. Shi X, Bai H, Wang J et al. Behavioral Assessment of Sensory, Motor, Emotion, and Cognition in Rodent Models of Intracerebral Hemorrhage. Front Neurol. 2021;12:667511

37. Lubrich C, Giesler P, Kipp M. Motor Behavioral Deficits in the Cuprizone Model: Validity of the Rotarod Test Paradigm. International journal of molecular sciences. 2022;23(19)

38. Fujimoto ST, Longhi L, Saatman KE et al. Motor and cognitive function evaluation following experimental traumatic brain injury. Neurosci Biobehav Rev. 2004;28(4):365–78

39. Lee HJ, Kim KS, Kim EJ et al. Brain transplantation of immortalized human neural stem cells promotes functional recovery in mouse intracerebral hemorrhage stroke model. *Stem cells (Dayton*, Ohio). 2007;25(5):1204–12

40. SUNDBERG RD, BROMAN H. The application of the Prussian blue stain to previously stained films of blood and bone marrow. Blood. 1955;10(2):160–6

41. Burghardt I, Tritschler F, Opitz CA et al. Pirfenidone inhibits TGF-beta expression in malignant glioma cells. Biochem Biophys Res Commun. 2007;354(2):542–7

42. Sun Y, Zhang Y, Chi P. Pirfenidone suppresses TGF-β1-induced human intestinal fibroblasts activities by regulating proliferation and apoptosis via the inhibition of the Smad and PI3K/AKT signaling pathway. Mol Med Rep. 2018;18(4):3907–3913

43. Hao H, Gabbiani G, Bochaton-Piallat M. Arterial smooth muscle cell heterogeneity: implications for atherosclerosis and restenosis development. Arteriosclerosis, thrombosis, and vascular biology. 2003;23(9):1510–20

44. Charidimou A, Boulouis G, Gurol ME et al. Emerging concepts in sporadic cerebral amyloid angiopathy. Brain : a journal of neurology. 2017;140(7):1829–1850

45. Cordonnier C, Leys D, Dumont F et al. What are the causes of pre-existing dementia in patients with intracerebral haemorrhages? Brain : a journal of neurology. 2010;133(11):3281–9

46. de Laat KF, van Norden AGW, Gons RAR et al. Gait in elderly with cerebral small vessel disease. Stroke. 2010;41(8):1652–8

47. Sharma B, Wang M, McCreary CR, Camicioli R, Smith EE. Gait and falls in cerebral small vessel disease: a systematic review and meta-analysis. Age Ageing. 2023;52(3)

48. Blevins BL, Vinters HV, Love S et al. Brain arteriolosclerosis. Acta Neuropathol. 2021;141(1):1–24

49. Pantoni L. Cerebral small vessel disease: from pathogenesis and clinical characteristics to therapeutic challenges. The Lancet. Neurology. 2010;9(7):689–701

50. Bullmore E, Sporns O. The economy of brain network organization. Nature reviews. Neuroscience. 2012;13(5):336–49

51. Hill RA, Tong L, Yuan P et al. Regional Blood Flow in the Normal and Ischemic Brain Is Controlled by Arteriolar Smooth Muscle Cell Contractility and Not by Capillary Pericytes. Neuron. 2015;87(1):95–110

52. Nishimura N, Schaffer CB, Friedman B, Lyden PD, Kleinfeld D. Penetrating arterioles are a bottleneck in the perfusion of neocortex. Proc Natl Acad Sci U S A. 2007;104(1):365–70

53. Frösen J, Joutel A. Smooth muscle cells of intracranial vessels: from development to disease. Cardiovasc Res. 2018;114(4):501–512

54. Yang Z, Wang J, Zheng T, Altura BT, Altura BM. Importance of extracellular Ca2+ and intracellular Ca2+ release in ethanol-induced contraction of cerebral arterial smooth muscle. *Alcohol (Fayetteville*, N.Y.). 2001;24(3):145–53

55. Liu P, Xi Q, Ahmed A, Jaggar JH, Dopico AM. Essential role for smooth muscle BK channels in alcohol-induced cerebrovascular constriction. Proc Natl Acad Sci U S A. 2004;101(52):18217–22

56. Altura BM, Zhang A, Cheng TP, Altura BT. Alcohols induce rapid depletion of intracellular free Mg2+ in cerebral vascular muscle cells: relation to chain length and partition coefficient. *Alcohol (Fayetteville*, N.Y.). 1995;12(3):247–50

57. Holm A, Heumann T, Augustin HG. Microvascular Mural Cell Organotypic Heterogeneity and Functional Plasticity. Trends Cell Biol. 2018;28(4):302–316

58. Buckwalter MS, Wyss-Coray T. Modelling neuroinflammatory phenotypes in vivo. J Neuroinflammation. 2004;1(1):10

59. Wyss-Coray T, Lin C, Sanan DA, Mucke L, Masliah E. Chronic overproduction of transforming growth factor-beta1 by astrocytes promotes Alzheimer’s disease-like microvascular degeneration in transgenic mice. The American journal of pathology. 2000;156(1):139–50

60. Wyss-Coray T, Masliah E, Mallory M et al. Amyloidogenic role of cytokine TGF-beta1 in transgenic mice and in Alzheimer’s disease. Nature. 1997;389(6651):603-6

61. Bozkurt I, Ozturk Y, Guney G et al. Effects of pirfenidone on experimental head injury in rats. Int J Clin Exp Pathol. 2022;15(1):20–28

62. Yang L, Wang F, Han H et al. Functionalized graphene oxide as a drug carrier for loading pirfenidone in treatment of subarachnoid hemorrhage. Colloids and surfaces. B, Biointerfaces. 2015;129:21–9

